# Tissue-specific transcriptomes reveal mechanisms of microbiome regulation in an ancient fish

**DOI:** 10.1101/2022.10.12.511976

**Authors:** Matt J. Thorstensen, Alyssa M. Weinrauch, William S. Bugg, Ken M. Jeffries, W. Gary Anderson

## Abstract

The lake sturgeon (*Acipenser fulvescens*) is an ancient, octoploid fish faced with conservation challenges across its range in North America but a lack of genomic resources has hindered molecular research in the species. To support such research we aimed to provide a transcriptomic database from 13 tissues: brain, esophagus, gill, head kidney, heart, white muscle, liver, glandular stomach, muscular stomach, anterior intestine, pyloric cecum, spiral valve, and rectum. The transcriptomes for each tissue were sequenced and assembled individually from a mean of 98.3 million (±38.9 million std. dev.) reads each. In addition, an overall transcriptome was assembled and annotated with all data used for each tissue-specific transcriptome. All assembled transcriptomes and their annotations were made publicly available as a scientific resource. The non-gut transcriptomes provide important resources for many research avenues, however, the gut represents a compartmentalized organ system with compartmentalized functions and the sequenced gut tissues were from each of these portions. Therefore, we focused our analysis on mRNA transcribed in different tissues of the gut and explored evidence of microbiome regulation. Gene set enrichment analyses were used to reveal the presence of photoperiod and circadian-related transcripts in the pyloric caecum, which may support periodicity in lake sturgeon digestion. Similar analyses were used to identify different types of innate immune regulation across the gut, while analyses of unique transcripts annotated to microbes revealed heterogeneous genera and genes among different gut tissues. The present results provide a scientific resource and information about the mechanisms of compartmentalized function across gut tissues in a phylogenetically ancient vertebrate.

## Introduction

The lake sturgeon (*Acipenser fulvescens*) is an octoploid, ancient fish with conservation challenges across its range in North America (1). Molecular resources for lake sturgeon can thus support wide-ranging research questions about fundamental biology relevant to their conservation. However, such research has been hampered by the limited molecular resources available for studying the species. While a microsatellite panel and genotyping by sequencing have been used for population genetic research (2–5), microsatellites are not as informative as reduced representation sequencing for individual genotype information, and may miss patterns of admixture and hierarchical structure (6,7). Moreover, reference-free reduced representation sequencing is more vulnerable to stochasticity in results than reference-based approaches (i.e., with a reference genome or transcriptome) due to the bioinformatics pipelines used (8). Sequencing resources such as reference transcriptomes or a well-annotated genome would thus enable more thorough molecular research, but the lack of sequence data for some species has complicated the development of new assays for research on stress responses. While some work has been done using specific primers developed to assay mRNA abundance in the species (9–11), the lack of a publicly available lake sturgeon transcriptome and genome has hindered molecular physiology and environmental DNA work (12).

Furthermore, the early divergence of sturgeons (13,14) make representative species such as lake sturgeon useful for studying questions about vertebrate evolution. For example, the pyloric caecum was first studied by Aristotle, who hypothesized about storage, fermentation, and digestive functions and caeca in fish digestive tracts were then later determined to increase gut surface area for digestion and absorption (15,16). Sturgeons represent the first evolutionary appearance of fused caeca with increased surface area (17,18), making these fish an important group for understanding the evolution of vertebrate digestive organs and function. An important caveat is that the presence and function of the lake sturgeon pyloric caecum should not be viewed as a basal state for Actinopterygii given that evolution in certain genes has been observed in other sturgeons and paddlefishes, and has presumably occurred in lake sturgeon as well (19–21). Nevertheless, the lake sturgeon pyloric caecum can be used as a representative tissue to study the evolution of vertebrate digestion (22).

Alternatively, the lake sturgeon may be useful for studying vertebrate digestion from a whole-organism perspective, as gut microbiomes have been described in a wide variety of organisms, including insects, fishes, and humans (23–26). The lake sturgeon microbiome has been linked to its physiological state, providing evidence for host-microbe interactions (27–29). However, regulation of gut microbiota across different gut tissues has been well-characterized in only a few species, mostly humans and lab mice (26,30–32). With messenger RNA sequencing, nearly all mRNA transcripts from a tissue can be assembled and annotated regardless of species. The genus of a given transcript annotation can be inferred by using taxonomic information from transcriptome annotations—even if that genus is within Bacteria or Archaebacteria. Therefore, RNA sequencing in the lake sturgeon can be used to study gut microbiome heterogeneity and regulation, with implications for the evolution of gut microbiome regulation across vertebrates. While the community structure of the microbiome is heavily influenced by environmental factors (33), the hypothetical presence of heterogeneity in genera and genes in the microbial community across the lake sturgeon gut would suggest the presence of tissue-specific microbial regulatory mechanisms. Moreover, fish may have been the first group of microbiome hosts to evolve an innate capacity for microbiome regulation (33). Ancient bony fish such as the lake sturgeon are thus valuable for studying the dynamics between host and microbiome.

While assembling a genome for a polyploid fish involves extensive chromatin and long-read DNA sequencing (20), assembling a transcriptome is a more tractable task. Such transcriptomes enable in-depth analyses of molecular physiology, such as in population-specific thermal stress responses (34–36). While RNA-seq and transcriptome-based approaches are less commonly used for population genetics than DNA-based approaches, single nucleotide polymorphisms in RNA can be used to investigate population structure and signatures of selection (37–41). Transcriptomes would also enable investigations into evolution, both through descriptions of gene expression evolution (42,43) and by phylogenetic analyses of mutations (44,45). Therefore, transcriptome assembly, annotation, and dissemination enables broad research questions in physiology and genetics.

Multi-tissue and tissue-specific approaches to transcriptome assembly allow for more systematic and in-depth analysis than typical transcriptomics, which may use one tissue for more focused investigations. For example, tissue-specific analyses revealed specialization in tissues and stages in cell division, photosynthesis, auxin transport, stress responses, and secondary metabolism in the tomato (*Solanum pimpinellifolium*) (46). In a livebearing fish, *Poeciliopsis prolifica,* multiple tissues were used with RNA-seq data to investigate placental evolution, where the abundance of clusters of transcripts was associated with different tissues (47). In the Atlantic salmon (*Salmo salar*), a blood-specific transcriptome was compared to other tissue transcriptomes to identify genes and gene ontology terms unique to blood (48). Thus transcriptome assembly with multiple tissues would provide a stronger resource for molecular research than single-tissue approaches.

One concern about transcriptomics in the lake sturgeon is that assembly may be affected by the octoploid status of the species (1). For instance, in situations where a transcriptome must be assembled without a reference genome, polyploid species are vulnerable to homeologs and ambiguous but similar sequences that decrease accuracy in the final assembly (49). One solution is to create transcriptomes specific to different conditions that may isolate different gene isoforms (49), a strategy consistent with the benefits of a multi-tissue, tissue-specific approach. In addition, different transcriptome assembly programs had varying success at accurately assembling polyploid transcriptomes, and careful selection of an assembly program such as Trinity can at least partly address challenges introduced by high ploidies (49–52).

In this study, we assembled, annotated, and analyzed the transcriptomes of 13 tissues in the lake sturgeon, sequenced with short-read messenger RNA sequencing (i.e., Illumina). The tissues were: brain, esophagus, gill, head kidney, heart, white muscle, liver, glandular stomach, muscular stomach, anterior intestine, pyloric cecum, spiral valve, and rectum (Fig. 1). In addition, all data used to assemble each of the 13 tissue-specific transcriptomes was assembled and annotated as an overall transcriptome. All transcriptomes and annotations were made publicly available for use as a scientific resource (https://figshare.com/projects/Lake_Sturgeon_Transcriptomes/133143). The brain, gill, head kidney, heart, white muscle, and liver transcriptomes have broad potential for studying many aspects of sturgeon biology. However, among the tissue-specific transcriptomes analyzed, the gut tissue transcriptomes represent components of an organ system with compartmentalized functions throughout (18). Because the 7 gut tissue transcriptomes were the major anatomically distinct regions of the lake sturgeon gut, these transcriptomes were used in more focused analyses and discussed in greater detail. Transcriptome annotations were analyzed in two ways: an exploratory approach using gene ontology terms (53) to identify unexpected transcript presence within and among tissues, and a guided approach informed by prior knowledge, where we used the lake sturgeon as a representative ancient fish to investigate different facets of vertebrate digestion. We focused on the pyloric caecum because the tissue first appears in sturgeons in terms of increased surface area for digestion and absorption (16,17). In addition, overall patterns of biological processes such as immune regulation across the gut and signatures of the lake sturgeon microbiome along different gut tissues were investigated. We observed circadian rhythm genes in the pyloric caecum, various types of innate immune regulation across gut tissues, and heterogeneity of bacteria and archaea associated transcripts in the lake sturgeon gut.

**Figure 1.**
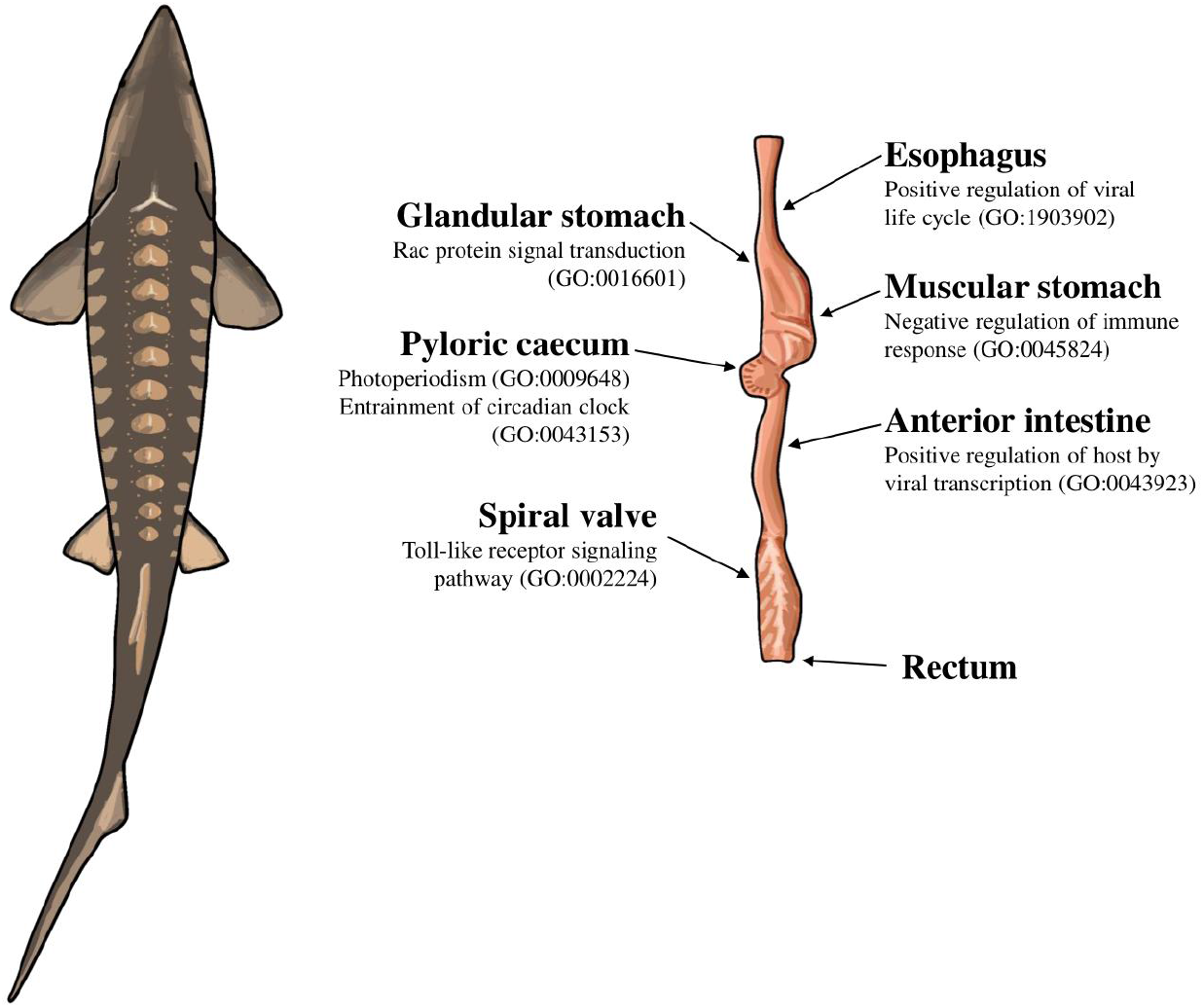
Illustration of a lake sturgeon (*Acipenser fulvescens*) and the gut tissues used for transcriptome assemblies in the present study. Beneath most gut tissues are representative, significant (*q* < 0.05), gene ontology terms unique to the tissue identified with EnrichR. The gene ontology terms present in the esophagus, glandular stomach, muscular stomach, anterior intestine, and spiral valve represent possible innate immune system processes specific to each gut tissue in the present transcriptomes. The gene ontology terms present in the pyloric caecum were processes related to circadian rhythms, unique to the tissue among the transcriptomes analyzed.

## Methods

### Sampling and Sequencing

Lake sturgeon of approximately 2 years old and unknown sex were sampled haphazardly from holding tanks and euthanized with an overdose of buffered tricaine methanesulfonate solution (MS-222; Sigma-Aldrich). Tissue from each of the gill, liver, brain, head kidney, white muscle, and heart were extracted from the sturgeon, stored in *RNAlater* (Thermo Fisher), and the PureLink RNA Mini Kit (Thermo Fisher) was used for RNA extractions following manufacturer protocols. By contrast, gut samples (esophagus, glandular stomach, muscular stomach, anterior intestine, pyloric caecum, spiral valve, and rectum) were immediately placed in Trizol and extracted the same day following the manufacturer’s protocol (Thermo Fisher) to limit RNA degradation that can be associated with digestive enzymes and gut bacteria naturally present in the tissues (54). For all tissues, equal amounts of RNA were pooled from *n*=3 fish for sequencing.

Total RNA was sent to the Centre d’expertise et de services Génome Québec, Montreal, Quebec (http://gqinnovationcenter.com), where 250 nanograms of total RNA per tissue was used with the NEBNext Poly(A) Magnetic Isolation Module (New England BioLabs). Stranded cDNA libraries were created with the NEBNext Ultra II Directional RNA Library Prep Kit for Illumina (New England Biolabs). Fish were sequenced for 100 base pair reads on one lane of a NovaSeq 6000 (Illumina). A mean of 98.3 million (± 38.9 million standard deviation (s.d.)) reads were sequenced for each tissue (Table 1).

**Table 1.**
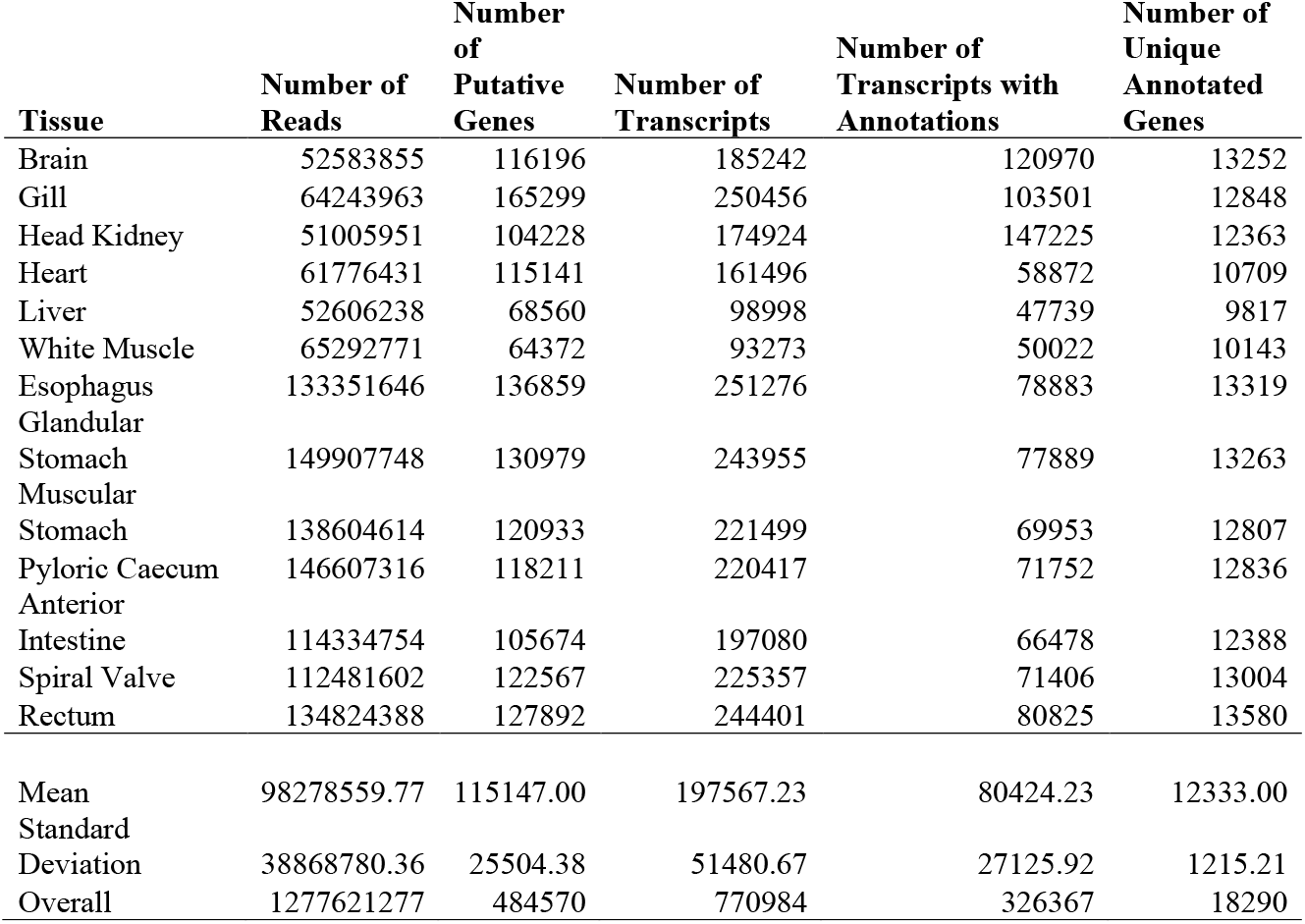
Assembly and annotation statistics for each of 13 tissue-specific transcriptomes of the lake sturgeon (*Acipenser fulvescens*). Number of reads represents the number of short read mRNA sequences used in assembly, number of putative genes refers to the number of gene models assembled in Trinity, and number of transcripts represents the total number of transcripts identified. The numbers of transcripts with annotations and number of unique annotated genes refers to annotations performed with Trinotate and associated programs. Means and standard deviations among all 13 transcriptomes are reported at the bottom of the table. The tissue-specific transcriptomes are labeled by tissue, while the transcriptome labeled ‘Overall’ refers to an assembly that included all data used to make the 13 tissue-specific transcriptomes.

### Transcriptome Assembly and Annotation

Trinity was used for transcriptome assembly (52,55), while Trinotate was used for transcriptome annotation (56–62). Both Trinity and Trinotate were used on mRNA sequencing data from each tissue separately as tissue-specific transcriptomes, and with all data from the 13 tissues as an overall transcriptome. Trinity was used for its effectiveness at assembling polyploid genomes (49). Transcriptome annotations were filtered for transcripts with E-values < 1×10^-6^ and bit scores > 50 (63). BUSCO v5.1.2 was used to assess transcriptome completeness with respect to the Actinopterygii odb10 dataset (64). The fishualize v0.2.3 package in the statistical computing environment R v1.1.2 was used to visualize results (65,66). To assess divergence in terms of mutation distance, Mash v1.1 was used to make pairwise comparisons between each transcriptome (67). Because evolutionary distance is not expected among transcriptomes from lake sturgeon sampled from a single population, the distances measured thus represent isoforms and paralogs among the gene models in the assembled transcriptomes.

### Annotation Analyses

The statistical computing environment R v1.1.2 and R package Tidyverse v1.3.1 were used throughout functional analyses of lake sturgeon transcriptomes (66,68). A gene set enrichment analyses was used to identify gene ontology terms for each transcriptome using enrichR v3.0 with the Biological Process 2021, Molecular Function 2021, and Cellular Component 2021 databases (53). Only gene ontology terms with a false discovery rate (*q*) < 0.05 were retained as significantly enriched. The R package UpSetR v1.4.0 was used to assess uniqueness of gene ontology terms among tissues (69). This assessment of uniqueness among tissues was repeated in an analysis excluding peripheral tissues and specific to the esophagus, glandular stomach, muscular stomach, pyloric caecum, anterior intestine, spiral valve, and rectum to identify patterns specific to lake sturgeon gut tissues.

Principal components analysis (PCA) was used to visualize differentiation among different tissues with respect to present genes and gene ontology terms. The overall transcriptome was excluded from PCA to explore variance among the tissue-specific transcriptomes. A PCA with *prcomp* in R was used on a table of presence or absence of gene names within each of the 13 tissues, along with separate PCAs for each of the Biological Process 2021, Molecular Function 2021, and Cellular Component 2021 gene ontology databases. These PCAs were used to visualize differentiation among the different transcriptomes. The overall transcriptome was generally excluded from annotation comparisons as analyses of uniqueness among tissues focused on the set of the tissue-specific transcriptomes. However, the number of unique genes was assessed in the overall transcriptome to identify genes that were potentially missing or un-annotated from each tissue-specific transcriptome, but resolved with all data used in one assembly. The genes unique to the overall transcriptome were analyzed for gene ontology terms with the same databases as the tissue-specific transcriptomes.

### Microbial Analyses

The lake sturgeon microbiome was investigated by removing transcripts annotated to eukaryotes or viruses from the transcriptomes of all 13 tissues. Therefore, only transcripts annotated to bacteria or archaea remained from each transcriptome. A gene set enrichment analysis was performed on the microbe-annotated transcripts, but no significant gene ontology terms were identified. UpSet plots were used to identify uniqueness in annotated genes and microbial genera present among tissues.

## Results

### Transcriptome Assembly and Annotation

The mean number of putative genes from the Trinity assemblies was 115,147 (± 24,405 s.d.), while the mean number of transcripts was 197,567 (± 51,481 s.d.) (Table 1). After filtering out transcripts with annotations with bit scores < 50 and E-values > 1×10^-6^, the number of transcripts remaining with annotations among the tissue-specific transcriptomes was a mean of 76,319 (± 19,737 s.d.) representing a mean 12,350 (± 1,217 s.d.) unique genes with annotations (Table 1). 3,065 genes were identified in analysis of uniqueness that included the overall and tissue-specific transcriptomes, although uniqueness from gene sets for the tissue-specific transcriptomes were skewed downward (Supplementary Figure S1). Tissue-specific transcriptome completeness ranged from a minimum of 33.7% (heart) to a maximum of 82.2% (rectum) (overall mean 65.1% ± 15.7% s.d.) (Fig. 2). Divergence among tissue-specific transcriptomes, as assessed by Mash, was statistically significant in each pairwise comparison (*p* < 0.05), although distances between transcriptomes were greatest between gut tissues and the heart, liver, and white muscle tissues (Fig. 3A).

**Figure 2.**
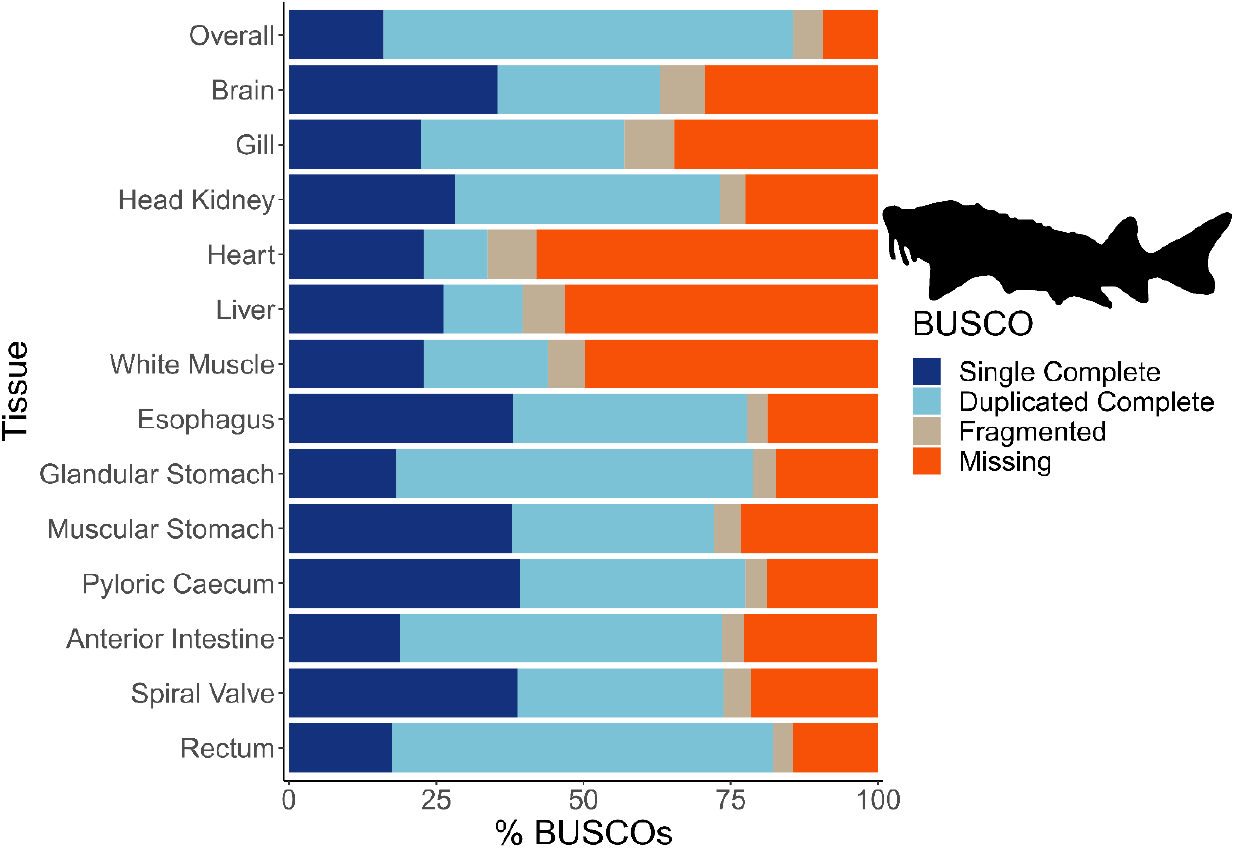
Transcriptome completeness assessed with BUSCO. Single complete represents orthologs that were present singly in a transcriptome, while duplicated complete represents orthologs duplicated in the transcriptome that matched the BUSCO profile. Fragmented orthologs were present in the transcriptomes, but not within the expected range of alignments in the BUSCO profile. Missing orthologs were those present in the BUSCO profile, but missing in the transcriptome completely. The tissue-specific transcriptomes are labeled by tissue, while the transcriptome labeled ‘Overall’ refers to an assembly that included all data from the 13 tissues. The BUSCO profile used in the present analysis was the Actinopterygii odb10 dataset. The lake sturgeon icon and colours used were from the fishualize package in R.

**Figure 3.**
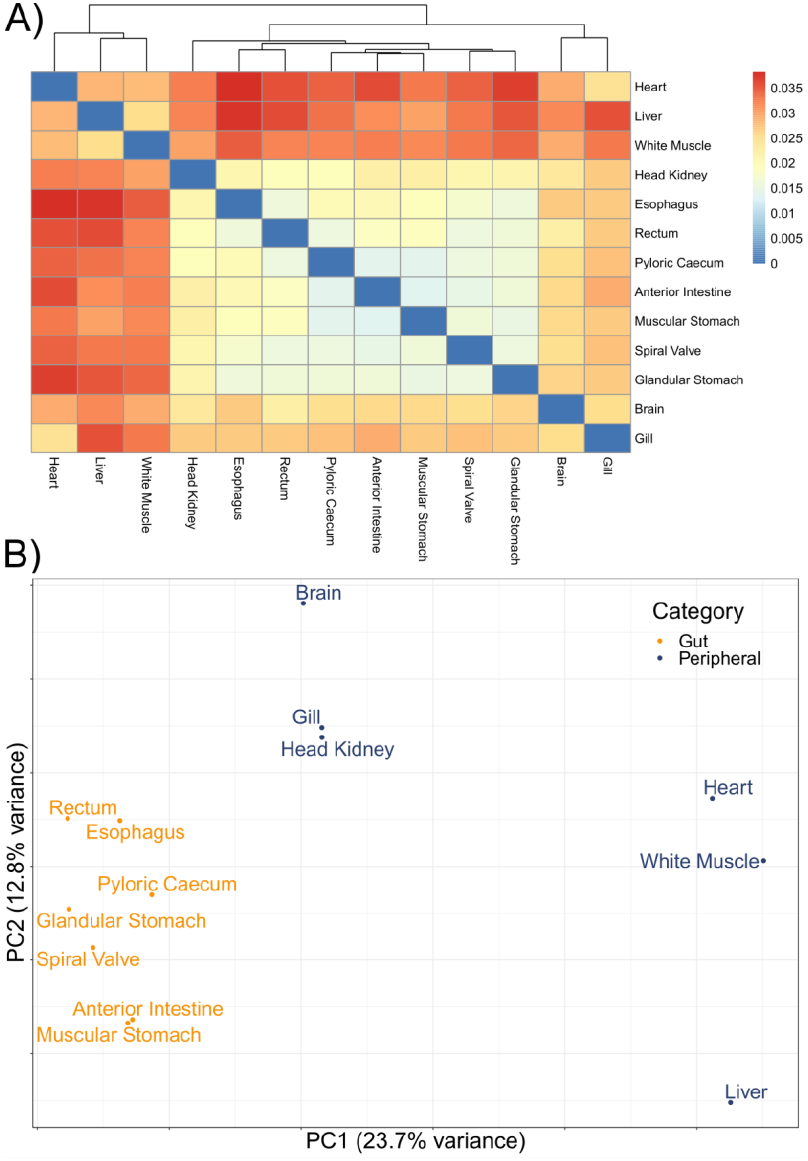
Divergence among 13 tissue-specific transcriptomes of the lake sturgeon (*Acipenser fulvescens*). A) is a heatmap of pairwise distances between transcriptomes assessed with Mash, where higher Mash distances correspond to greater evolutionary divergence between the transcriptomes. Because no evolutionary divergence is expected for transcriptomes from one population of one species, these distances represent isoforms and paralogs of gene models. Higher values indicate more divergence. B) is a principal components analysis (PCA) of present and absent genes in the 13 transcriptomes, performed with *prcomp* in R. Gut and peripheral tissues were distinguished for visualization, where gut tissues were the esophagus, glandular stomach, muscular stomach, anterior intestine, spiral valve, and rectum, while peripheral tissues were the brain, gill, head kidney, heart, white muscle, and liver. The distinction in colour between gut and peripheral tissues is only for visualization, and was not used to categorize data *a priori* in the PCA.

### Annotation Analyses

A mean of 832 (± 123 s.d.) biological process gene ontology terms were identified among the 13 tissue-specific transcriptomes (Table 2). Among biological process gene ontology terms, 367 were shared across all tissues, but a substantial number were also unique to individual tissues such as 101 gene ontology terms in the liver and 71 in the heart (Fig. 4; Supplementary Tables S1-S13). Qualitatively similar patterns of shared gene ontology terms among all tissues, with substantial numbers of terms unique to each tissue, were observed in the molecular function and cellular component databases (Supplementary Figures S2 & S3). A similar pattern of uniqueness was also present among biological process gene ontology terms in the gut tissues. Here, 464 terms were shared among all gut tissues, but smaller numbers of terms were unique to individual tissues, such as 59 unique to the anterior intestine and 46 unique to the pyloric caecum (Supplementary Figure S4). For molecular function and cellular component, a mean of 131 (± 19 s.d.) and 150 (± 21 s.d.) gene ontology terms were identified, respectively. Molecular function and cellular component gene ontology terms were qualitatively similar in patterns of uniqueness both when considering all 13 tissues (Supplementary Tables S1-S13) and among gut-only tissues (Supplementary Tables S14-S20). No significant gene ontology terms were identified from the overall transcriptome, possibly because the Fisher exact test used in enrichR was implemented for experimental designs, as opposed to surveys of gene presence (70). Five molecular function gene ontology terms were significant in a search of the 3,065 genes unique to the overall transcriptome among all transcriptomes analyzed, while no biological process or cellular component terms were significant. The significant molecular function gene ontgoloy terms were: peptide alpha-N-acetyltransferase activity (GO:0004596; combined score = 329), peptide N-acetyltransferase activity (GO:0034212; combined score = 255), phosphatidate phosphatase activity (GO:0008195; combined score = 251), lipid phosphatase activity (GO:0042577; combined score = 173), and lysine N-methyltransferase activity (GO:0016278; combined score = 38). PCAs revealed differentiation between the liver transcriptome and those from other tissues in each comparison, especially in a PCA of terms in the cellular components gene ontology database (Fig. 3B; Supplementary Figure S5). Gut tissues tended to cluster together compared to other tissues in each PCA, but because gut and peripheral tissues were extracted using separate protocols, some differentiation between the two groups of tissues may be a technical artifact.

**Figure 4.**
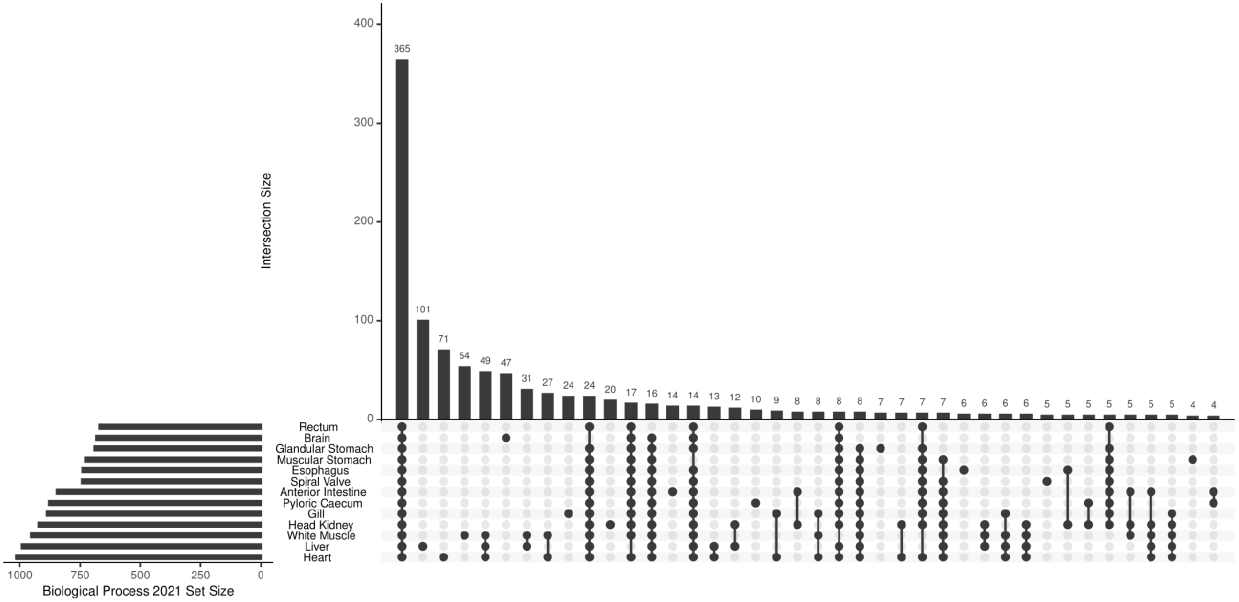
UpSet plot of shared and unique gene ontology (GO) terms from each of 13 tissue-specific transcriptomes of the lake sturgeon (*Acipenser fulvescens*). The GO terms presented here are from the Biological Process 2021 database, significant at *q* < 0.05. The R package UpSetR was used to visualize these data.

**Table 2.** Gene ontology (GO) terms and microbial information for each of 13 tissue-specific transcriptomes of the lake sturgeon (*Acipenser fulvescens*). GO terms were identified by using all unique annotated genes in a transcriptome with EnrichR. The Biological Process 2021, Molecular Function 2021, and Cellular Component 2021 databases were searched for the GO analyses, where only significant terms (*q* < 0.05) were retained for downstream analyses. Present microbial genera and genes were identified using annotation information from Trinotate.

| **Tissue** | **Biological**<br>**Process GO**<br>**Terms** | **Molecular**<br>**Function**<br>**GO Terms** | **Cellular**<br>**Component**<br>**GO Terms** | **Microbial**<br>**Genera** | **Microbial**<br>**Genes** |
| --- | --- | --- | --- | --- | --- |
| Brain | 684 | 111 | 152 | 25 | 31 |
| Gill | 888 | 145 | 149 | 24 | 33 |
| Head Kidney | 924 | 149 | 152 | 25 | 28 |
| Heart | 1017 | 148 | 174 | 16 | 22 |
| Liver | 992 | 164 | 183 | 21 | 32 |
| White Muscle | 954 | 152 | 186 | 12 | 16 |
| Esophagus | 742 | 131 | 138 | 56 | 105 |
| Glandular Stomach | 691 | 110 | 131 | 45 | 83 |
| Muscular Stomach | 729 | 117 | 135 | 75 | 177 |
| Pyloric Caecum | 879 | 126 | 157 | 30 | 55 |
| Anterior Intestine | 848 | 136 | 145 | 38 | 67 |
| Spiral Valve | 744 | 119 | 124 | 65 | 169 |
| Rectum | 671 | 98 | 118 | 47 | 84 |
| Mean | 827.92 | 131.23 | 149.54 | 36.85 | 69.38 |
| Standard Deviation | 118.55 | 18.93 | 20.51 | 18.77 | 51.43 |

In the pyloric caecum, the gene ontology terms photoperiodism (GO:0009648) and entrainment of circadian clock by photoperiod (GO:0043153) were uniquely present and related to periodicity (Supplementary Table S11). Patterns of tissue-specific immune regulation were uniquely present in several gut tissues. Rac protein signal transduction (GO: 0016601) was uniquely present in the glandular stomach and may also represent a part of the innate immune system with its role in neutrophil recruitment (Supplementary Table S8). Negative regulation of immune response (GO:0045824) and autophagy of peroxisomes (GO:0030242) were unique to the muscular stomach (Supplementary Table S9). Positive regulation of host by viral transcription (GO:0043923) was uniquely present in the anterior intestine (Supplementary Table S10). Toll-like receptor 9 signaling pathway (GO:0034162), toll-like receptor signaling pathway (GO:0002224), and cellular response to interleukin-12 (GO:0071349) were each uniquely present in the spiral valve, consistent with a role for the tissue in the innate immune system (Supplementary Table S12). Positive regulation of viral life cycle (GO:1903902) was uniquely present in the esophagus (Supplementary Table S2).

### Microbial Analyses

A mean of 38 (± 19 s.d.) bacterial and archaeal genera were observed among 13 transcriptomes. A mean of 73 (± 52 s.d.) genes were annotated to bacteria or archaea among the same 12 transcriptomes. Both microbial genera and annotated genes showed a pattern of high uniqueness in each tissue, although 8 genera and 7 genes were present among all tissues (Fig. 5; Supplementary Figure S6).

**Figure 5.**
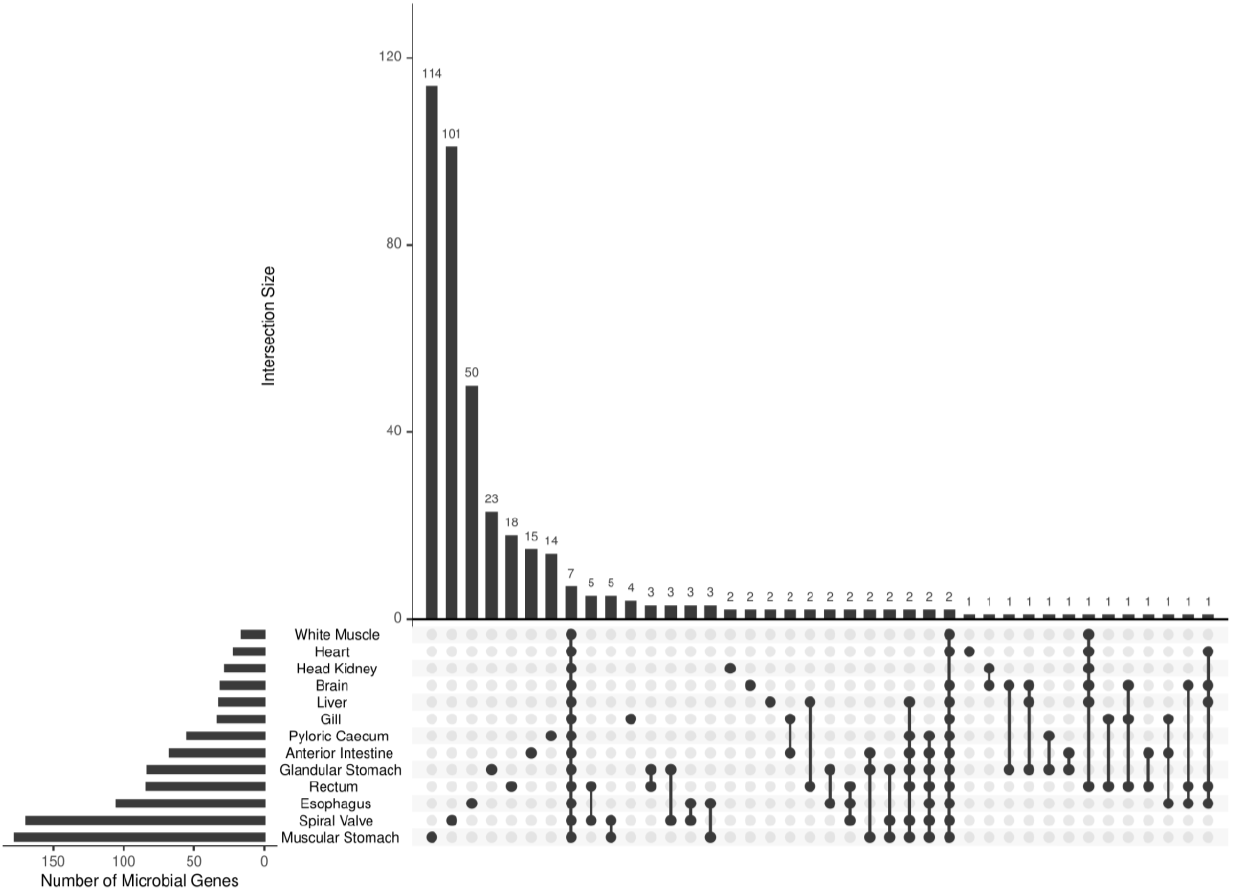
UpSet plot of shared and unique genes annotated to microbes (bacteria or archaea) from each of 13 tissue-specific transcriptomes of the lake sturgeon (*Acipenser fulvescens).* Annotations were performed with Trinotate, and bacterial or archaeal genes were identified by filtering for those groups among filtered transcriptome annotation reports.

## Discussion

### A Database of Transcriptomes

The 13 tissue-specific transcriptomes and one overall transcriptome presented in this study are a genomic resource publicly available for studying sturgeons. Transcriptomes are most commonly used for molecular physiology, and the gill transcriptome presented here has already been applied to study thermal stress between latitudinally separated populations of lake sturgeon (36). These transcriptomic resources may be used to characterize physiological responses to environmental conditions, which may in turn be used to inform conservation management (71). As lake sturgeon represent a species that exhibits extensive phenotypic plasticity (9,72,73), these transcriptomes also have potential for supporting fundamental research on the molecular basis of resilience to environmental change. Gene ontology terms revealed tissue-specific patterns in each of the transcriptomes presented here, such as 101 biological process terms unique to the liver and 71 unique to the heart. These terms thus represent transcriptional processes that would otherwise have been missing from the transcriptome database if only a single tissue was considered. Therefore, the 13 tissues we studied enable a broad range of analyses that would be otherwise intractable, allowing for in-depth assessments of shared and tissue-specific processes, along with genetic and physiological studies.

Among the 13 tissue-specific transcriptomes assembled, 7 were from the gut (esophagus, glandular stomach, muscular stomach, pyloric caecum, anterior intestine, spiral valve, and rectum), and 6 were from peripheral tissues (brain, gill, head kidney, heart, liver, and white muscle). The gut and peripheral tissue transcriptomes separated into different groups using two methods (PCA with present genes and metagenome distance estimation), although some separation between the two groups of tissues may be attributed to different RNA extraction methods used. Nevertheless, the distinction between the two groups is consistent with differences in physiological function. The peripheral tissue transcriptomes enable a variety of research questions on the lake sturgeon, such as liver and gill often used in work exploring the vertebrate stress response (74,75). Tissues such as the brain, heart, head kidney, and white muscle can be informative for developmental questions, with potential connections to nutrition and stress among other biological processes (74,76–79). Meanwhile, the 7 gut transcriptomes represent the major anatomically distinct regions of the gut. Because the gut encompasses an organ system with distinct compartmentalization of function (18), the 7 gut transcriptomes provide an opportunity to study distinct digestion-related mechanisms. Gut transcriptomics was predicted to accelerate research on intestinal pathogen responses, dietary manipulations, and osmoregulatory challenges (26), and the present data contribute to a longstanding body of work investigating physiological mechanisms of the vertebrate gut (15,16). We thus provide all 13 tissue-specific transcriptomes and one overall transcriptome as a scientific resource from this study, but focus on discussing observations among the gut tissues.

### Circadian Rhythm Transcripts in the Pyloric Caecum

The pyloric caecum is a tissue of interest because it is absent in Agnatha and Chondrichthyes, but is present in Actinopterygii (17,18,22). Given sturgeon’s status as an ancient actinopterygian, sister to the rest of the clade (13), they represent an early evolutionary appearance of the pyloric caecum. Analyses of tissue-specific transcriptome annotations revealed notable patterns of transcript presence within pyloric caecum, which may have implications for different mechanisms of lake sturgeon digestion. For instance, gene ontology terms photoperiodism (GO:0009648) and entrainment of circadian clock by photoperiod (GO: 0043153) were unique to the pyloric caecum. In addition, among the core clock genes of *clock, bmal1, per* (*1, 2,* and *3*), and *cry* (*1* and *2*), all expressed in teleost pineal organs (80), *clock, cry1*, and *cry2* were present in the lake sturgeon pyloric caecum. *Cry1* and *cry2*, which are photoreceptors with important roles in circadian rhythms, were also transcribed in brain, liver, heart, retina, muscle, spleen, gill, and intestine of European seabass (*Dicentrarchus labrax*), where rhythmic expression was observed in the brain and liver (81). Notably, the clock-related gene ontology terms observed in the present data were unique to the pyloric caecum, but individual genes were present among other tissues such as the brain. Gene ontology terms used in the present analyses were filtered for significance from a Fisher exact test (70). Therefore, transcriptomes with annotated clock genes but without enriched gene ontology terms present represent those with too few genes within the clock-related gene ontology terms to be significant. The present results do not contradict prior work that identified clock genes in other tissues, but do provide novel findings of core clock genes in the pyloric caecum that may be related to feeding periodicity.

Feeding periodicity has been observed in numerous fish species (e.g., *Merlagius merlangus,* (82); *Limanda limanda,* (83)), across herbivores, detritovores, insectivores, zooplanktivores, and macrophyte feeders (84). Because feeding periodicity is phylogenetically and ecologically widespread among fishes, we predict that physiological digestive mechanisms may contribute to the phenomenon. A circadian rhythm in metabolic rate was observed in lake sturgeon exposed to a 12-hour light-dark cycle, where metabolism was highest at sunrise (85). The lake sturgeon used for the present study were also fed predictably, three times a day with a 12-hour light-dark cycle. Given the dual observations that feeding periodicity exists across fish species (84) and the presence of several core clock genes in the lake sturgeon pyloric caecum, we developed alternative hypotheses that may address underlying mechanisms of periodicity in the lake sturgeon, that may be applicable in other fishes (86).

First, we hypothesized that physiological mechanisms of digestion periodicity may be regulated by diel circadian clock rhythms in the pyloric caecum. Therefore, we predict diel fluctuations in transcript abundance of *clock, cry1,* and *cry2* along with other circadian rhythm-related genes only in the pyloric caecum of laboratory-held lake sturgeon consistent with feeding times and a 12-hour light-dark cycle. Specific roles for physiological mechanisms of digestion periodicity could include intestinal motility, intestinal function, innate immunity, microbiome regulation, or cell proliferation (87–94). Alternatively, we hypothesized that the circadian rhythm genes observed in the pyloric caecum may represent a part of a whole-gut circadian response wave consistent with a phenomenon hypothesized in lab mice (95). That is, a whole-gut circadian response may have been initiated at feeding and was observed by chance in the pyloric caecum by sampling individuals approximately 17-18 hours after feeding. Therefore, from this hypothesis we predict diel fluctuations of transcript abundance of *clock*, *cry1*, and *cry2* in the pyloric caecum, and other gut tissues. More posterior gut tissues, such as the spiral valve, may therefore show expression of the three predicted genes but chronologically later than the pyloric caecum. By contrast, the glandular and muscular stomachs may show evidence of this circadian response wave earlier in time than the pyloric caecum because of their position prior to the caecum in the gut. While the first hypothesis about physiological functions of digestion as regulated by the pyloric caecum focuses on one gut tissue, the hypotheses are not necessarily mutually exclusive. A circadian response wave may pass through different gut tissues, but the pyloric caecum may play key roles in downstream processes from the circadian wave. This circadian response wave may be entrained by food intake times if it is consistent with mammalian physiology (96). Therefore, a timepoint- and tissue-specific approach is needed to test this hypothesis.

### Microbial Observations

As lake sturgeon represent an early stage of Actinopterygian gut evolution, several observations in the present database of tissue-specific transcriptomes were notable. One example is the presence of transcripts related to innate immunity in the spiral valve. As gut tissues may be in contact with food and potential associated pathogens from the external environment, innate immunity and immune responses involved in digestion may help to protect the fish from food or environment related pathogens (97). Cartilaginous fishes have gut-associated lymphoid tissues in the spiral valve (98,99), consistent with the present observations of innate immune-related transcripts in the lake sturgeon spiral valve. Gene ontology terms related to toll-like receptor signaling pathways and cellular responses to interleukin were unique to the spiral valve in lake sturgeon, while terms related to innate immune function were also present in the muscular stomach, glandular stomach, and anterior intestine. The gene ontology term positive regulation of viral life cycle is not an immune response in itself, but its unique presence in the esophagus provides some evidence for the necessity of an innate immune response in other gut tissues. These heterogeneous signals of host-microbiome interactions along the gut are consistent with evidence of host-microbiome interactions from 16S rRNA and DNA sequencing from microbial samples and the lake sturgeon spiral valve (27).

The tissue-specific transcriptome database enabled analyses of bacterial and archaeal transcripts as well as genera across different gut tissues. Many of these microbial transcripts and genera were unique to different gut sections, thus we concluded that the microbiome is likely heterogenous across the lake sturgeon gut. A caveat is that the presence of microbial genera and transcripts in certain tissues may be attributed to contamination, such as in the brain, white muscle, and heart, along with transcripts or genera shared among all tissues (7 genes and 8 genera identified as shared among all 13 tissues). However, patterns of microbial presence were consistent with microbiome regulation and tissue-specific function for gut tissues. For example, among microbial genera unique to each tissue, the muscular stomach had the greatest number present (15), followed by the spiral valve (12), and the anterior intestine (6). A qualitatively similar pattern was found with transcripts of genes annotated to bacteria and archaea unique to each tissue, with the muscular stomach (114), spiral valve (101), esophagus (50), glandular stomach (23), rectum (18), and pyloric caecum (14) all supporting unique microbial communities. These results demonstrate that the greatest number of unique microbial genera and genes were identified in gut tissues as opposed to tissues outside of the gut. Therefore, the present results are consistent with a heterogeneous microbial community with tissue-specific mRNA transcription in the lake sturgeon gut.

Other work identified microbial community shifts in the lake sturgeon spiral valve in response to a failure to transition diets and with feeding cessation (29). Similarly, spiral valve microbiome community composition changed in response to exposure to common antibiotics, drugs, and chemicals used in lake sturgeon aquaculture (28). Therefore, gut microbiome community composition is dynamically connected to the physiological state of lake sturgeon (27). Because unique patterns of microbial genera and genes were found in both the spiral valve and other gut tissues in the present data, the analyses of gut transcriptomes demonstrate that host-microbiome interactions may occur along much of the lake sturgeon gut, and that the interactions may be spatially heterogeneous and specific to different gut tissues. Thus the transcriptomes used here may support work in the lake sturgeon that resolves spatially distinct mechanisms of host-microbiome interactions.

### Conclusions

In the present study, 13 tissue-specific transcriptomes and one overall transcriptome were presented as a resource for lake sturgeon research. Overlap of gene ontology terms was analyzed among tissues. While shared patterns indicated consistent transcriptomic functions among tissues, the presence of unique gene ontology terms showed that sequencing transcriptomes from multiple tissues enabled research questions that would otherwise be intractable. Moreover, the analysis of unique gene ontology terms among tissues revealed the presence of transcribed genes related to photoperiodicity in the pyloric caecum, an observation consistent with a role for periodicity in digestive physiology in an ancient fish. Transcripts involved in innate immune function were found in the spiral valve and other gut tissues, which provide evidence in support of a prior hypothesis about the emergence of innate immunity in the gut of cartilaginous fishes and are consistent with specialization in immune function across gut tissues. An analysis of genes annotated to bacteria and archaea indicated potentially heterogeneous microbiota and microbial functions along different gut tissues, consistent with specialization in immune function and microbiome regulation along the lake sturgeon gut. As lake sturgeon are representative of sturgeon and paddlefish’s status as ancient fishes, they constitute an early stage in the differentiation of several gut tissues. Studying this early stage in differentiation of the gut as an organ system with distinct functions provided insights into digestive function, immunity, and microbiome regulation. These results are both a resource for lake sturgeon research and provide information about the mechanisms of compartmentalized function across gut tissues.

## Ethics approval and consent to participate

Not applicable.

## Consent for publication

Not applicable.

## Data availability

The lake sturgeon transcriptomes and annotations are available on Figshare https://figshare.com/projects/Lake_Sturgeon_Transcriptomes/133143. Code used in analyses of the data in the present study are available on GitHub (https://github.com/BioMatt/lakesturgeon_transcriptomes).

## Acknowledgements

We thank Dr. Robert Syme and François Lefebvre from the Canadian Centre for Computational Genomics for their support with mRNA sequencing. Evelien de Greef illustrated the lake sturgeon and gut, and Hamza Amjad assisted with RNA extractions.

## Authors’ contributions

M.J.T. performed bioinformatics analyses on the data and wrote the initial draft of the manuscript, with assistance from A.M.W. and W.S.B. All authors edited the manuscript. In addition, all authors designed and conceived of the study. W.G.A and K.M.J. provided funding and supervision.

