## Supplementary Figure for "Tissue-specific transcriptomes reveal mechanisms of microbiome regulation in an ancient fish"

Supplementary Figure S1. Uniqueness of gene sets among all transcriptomes, including the overall and tissue-specific transcriptomes as visualized with UpSetR. While 3,065 genes are uniquely present in the overall transcriptome, measures of uniqueness for the tissue-specific transcriptome were biased by inclusion of the overall transcriptome that was made with data used to assemble all of the others. Therefore, analyses of uniqueness among tissues focused on the set of the tissue-specific transcriptomes.

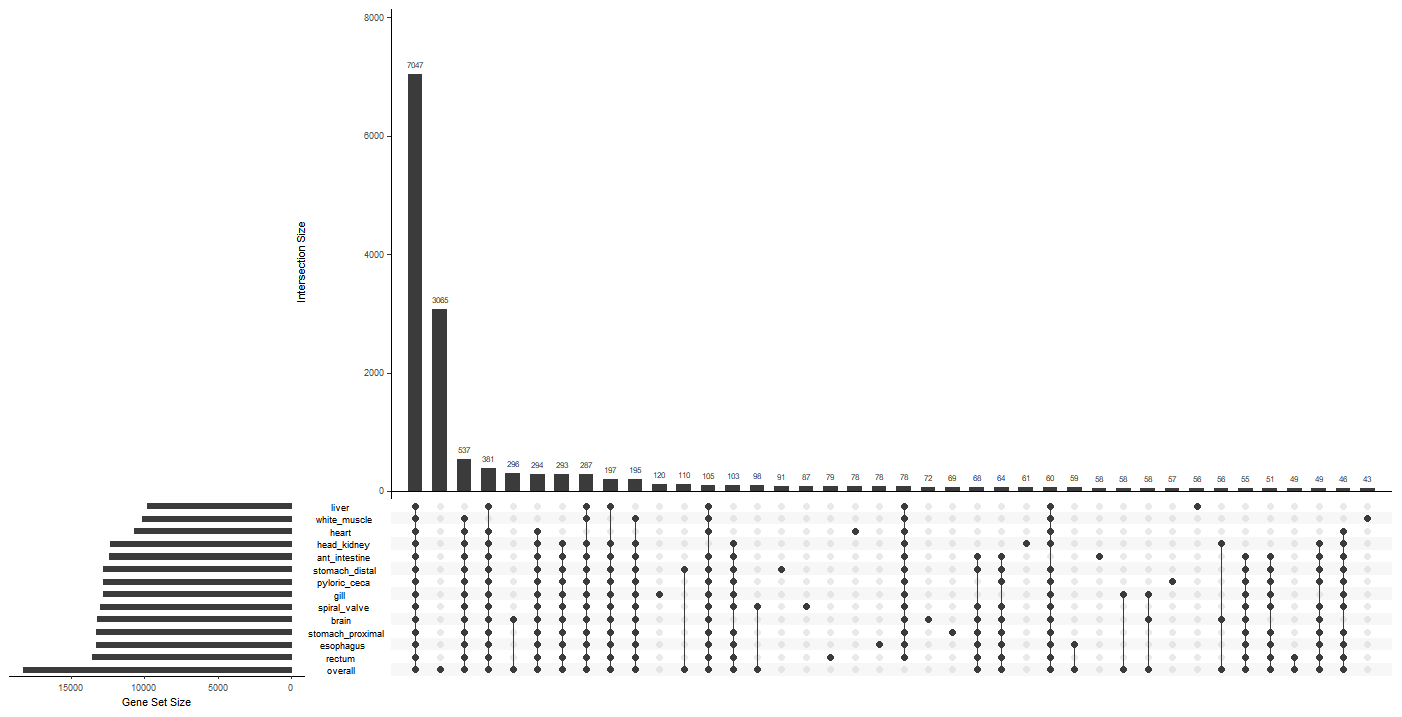

Supplementary Figure S2. Uniqueness of gene ontology terms from the Molecular Function 2021 database among 13 tissue-specific transcriptomes, as assessed with EnrichR and visualized with UpSetR.

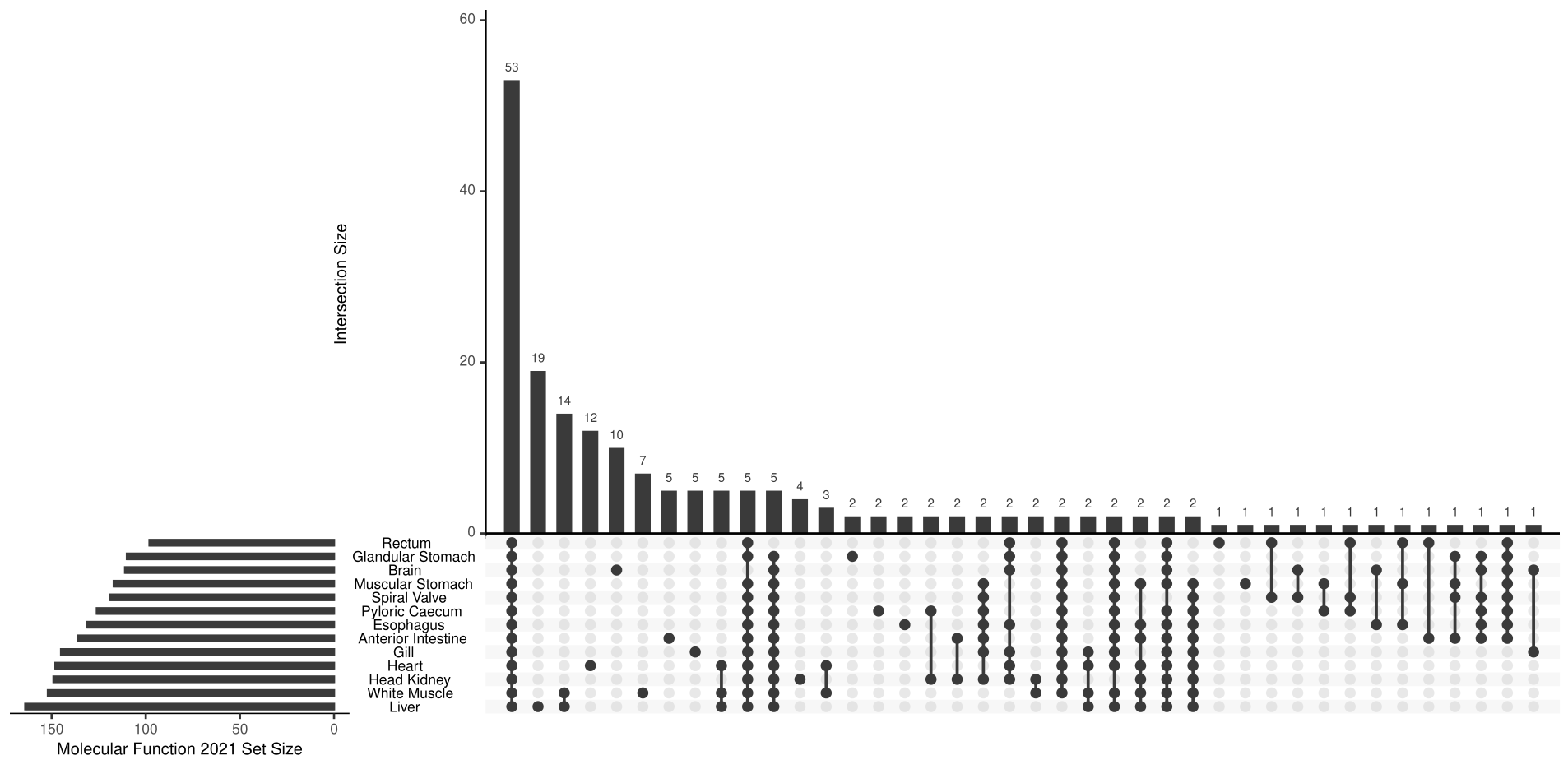

Supplementary Figure S3. Uniqueness of gene ontology terms from the Cellular Component 2021 database among 13 tissue-specific transcriptomes, as assessed with EnrichR and visualized with UpSetR.

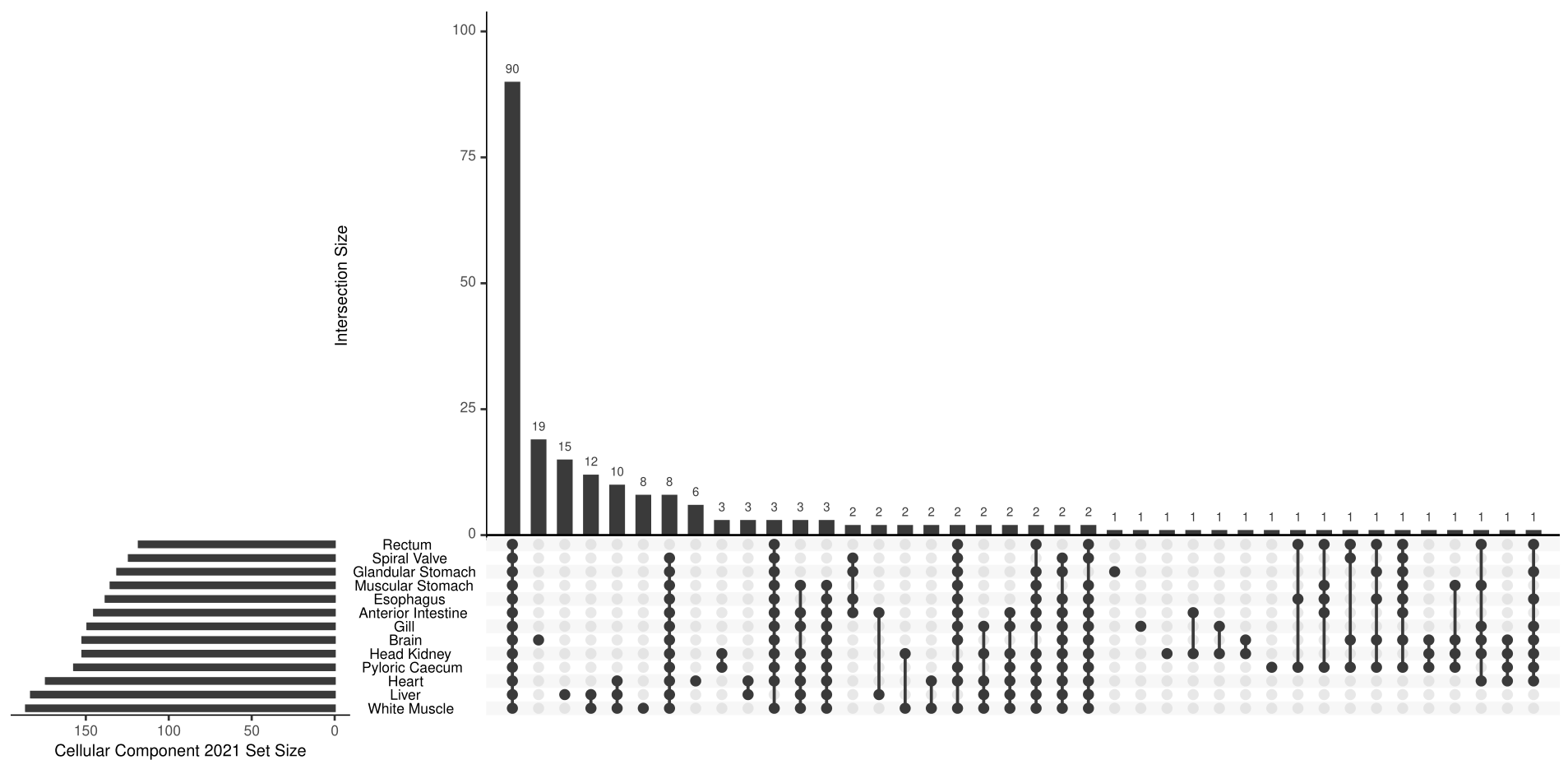

Supplementary Figure S4. Uniqueness of gene ontology terms from the Biological Process 2021 database among 7 gut tissue-specific transcriptomes, as assessed with EnrichR and visualized with UpSetR.

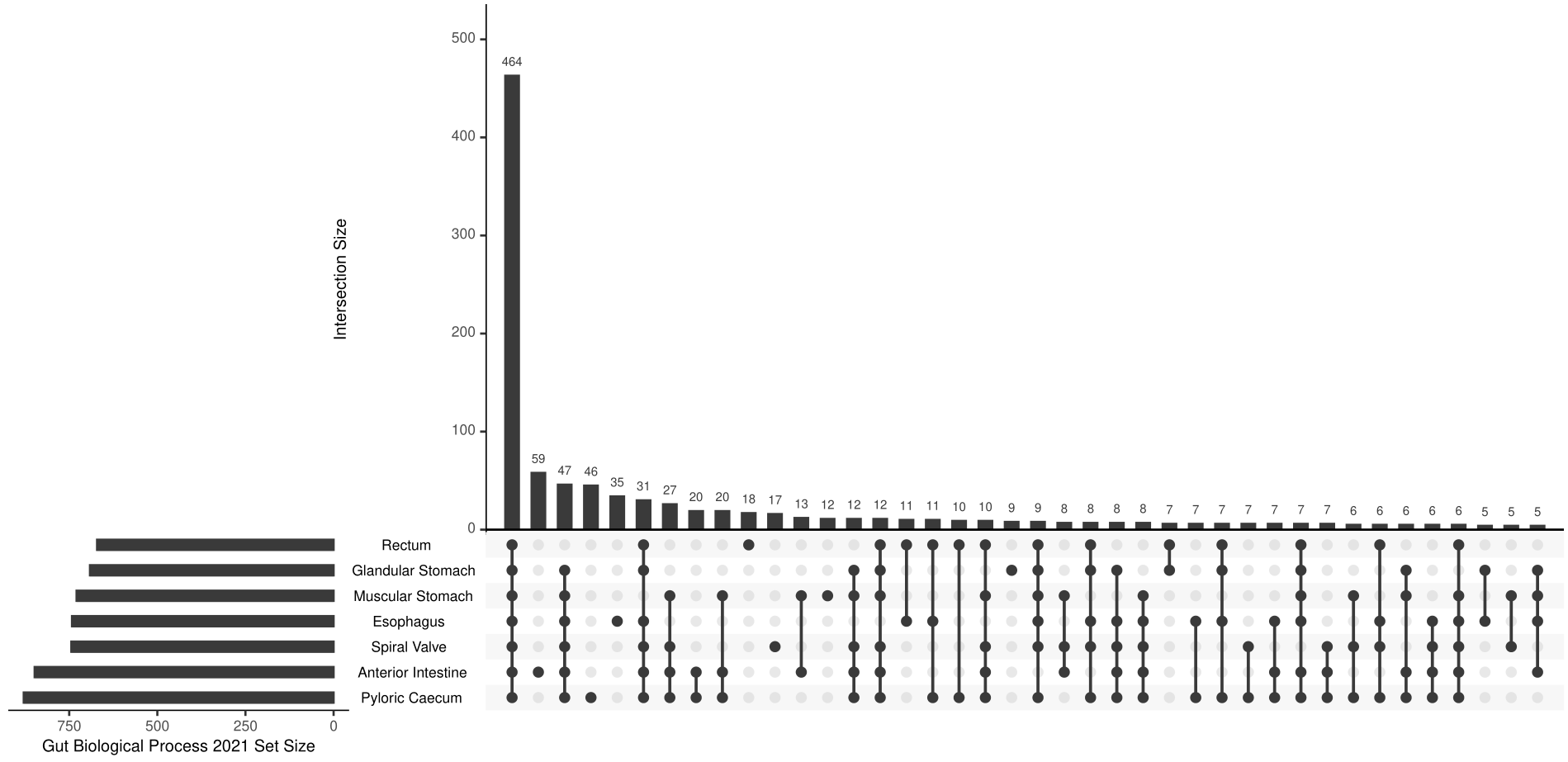

Supplementary Figure S5. Principal components analyses of present gene ontology terms among 13 tissue-specific transcriptomes as assessed with EnrichR and visualized with UpSetR. A) represents an analysis of terms from the biological process 2021 database, B) an analysis of terms from the molecular function 2021 database, and C) an analysis of terms from the cellular component 2021 database.

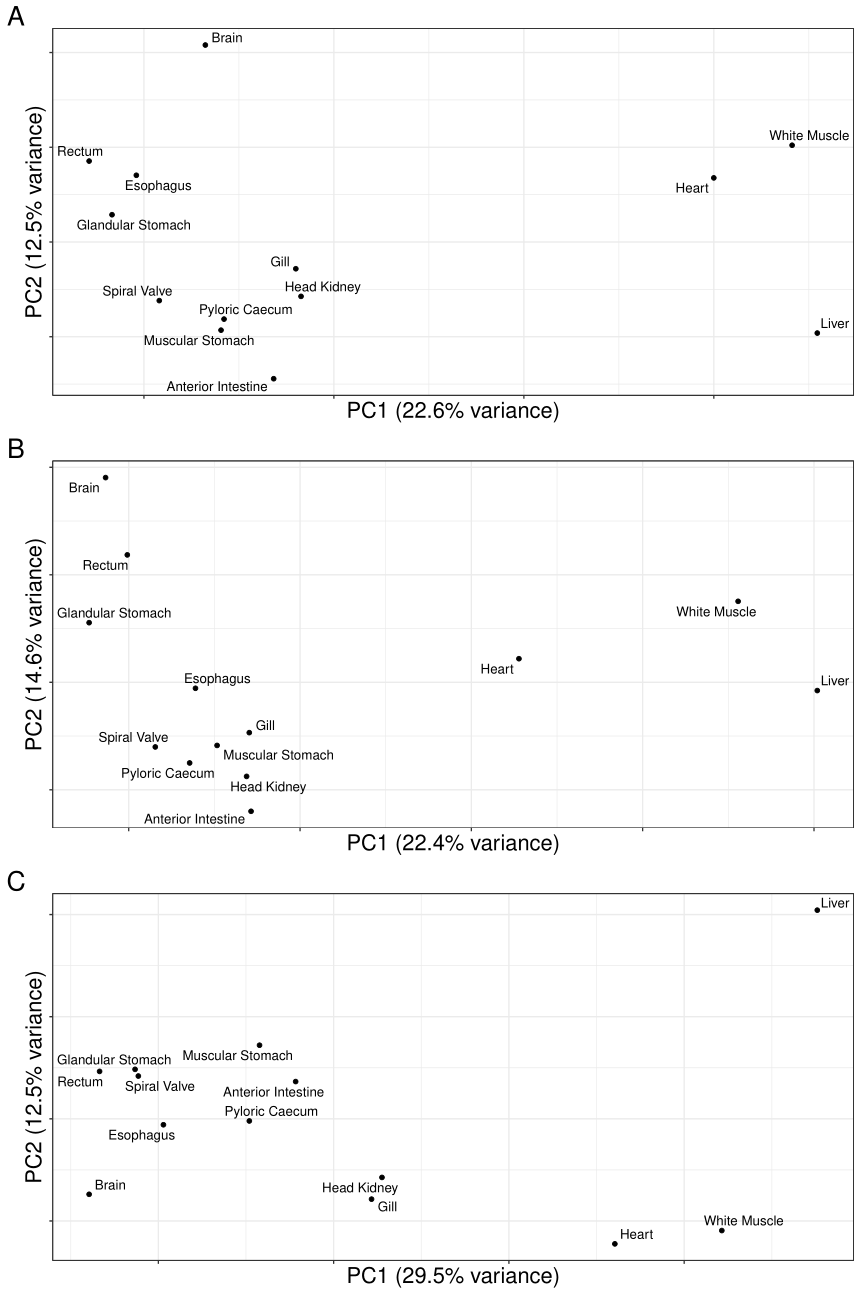

Supplementary Figure S6. Uniqueness of microbial (bacteria, viruses, and archaea) genera among 13 tissue-specific transcriptomes as visuzalized by UpSetR.
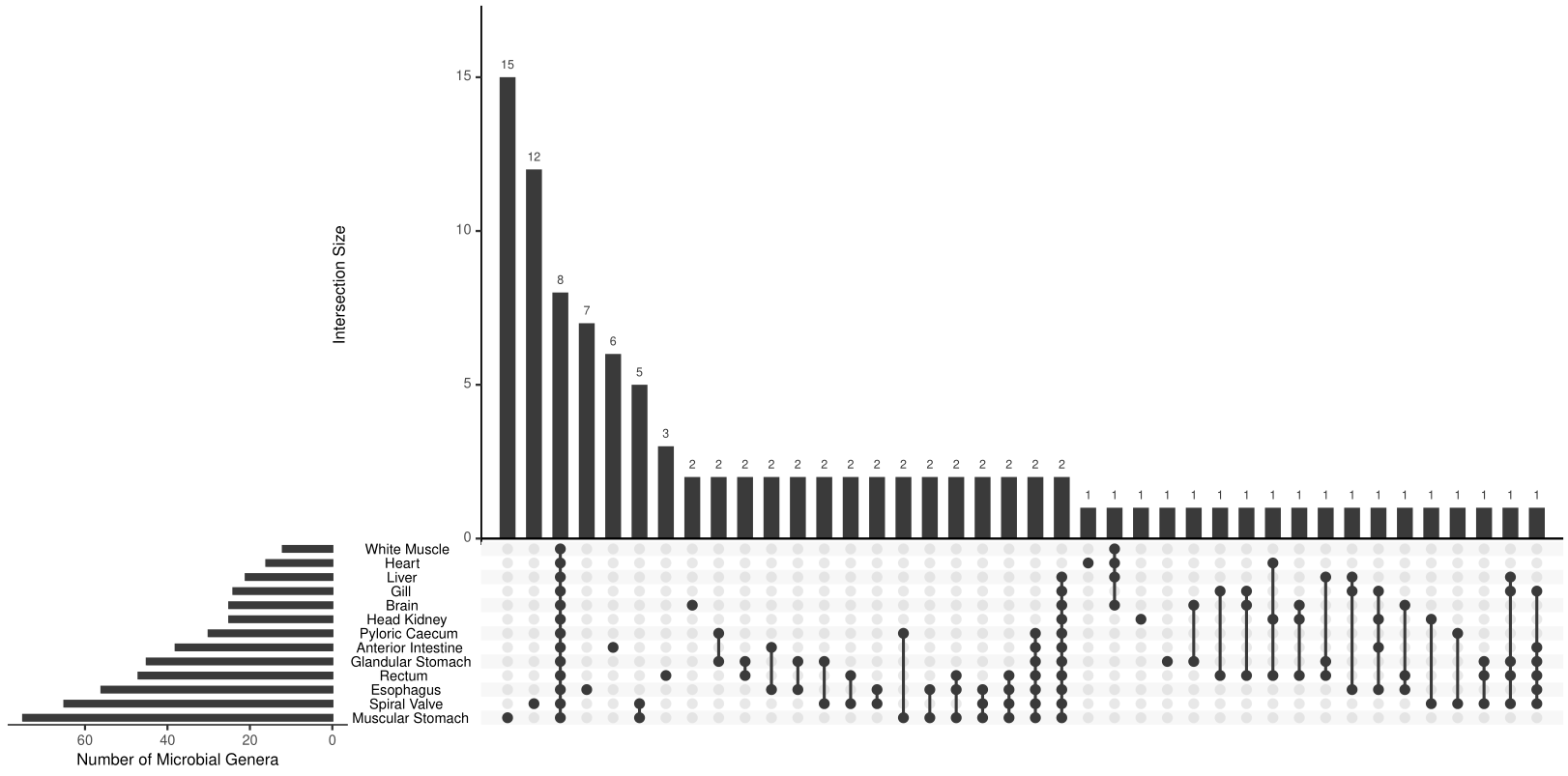
